# Inconsistent relationships detected between seed size, shape, and persistence for different plant functional groups in the Pannonian flora

**DOI:** 10.1101/2025.05.05.652257

**Authors:** Viktória Törő-Szijgyártó, Péter Török, Katalin Tóth, Hajnalka Málik-Roffa, Luis Roberto Guallichico Suntaxi, Szilvia Madar, Gergely Kovacsics-Vári, Andrea McIntosh-Buday, Patricia Díaz Cando, Judit Sonkoly

**Affiliations:** Department of Ecology, University of Debrecen, 1 Egyetem sqr., 4032 Debrecen, Hungary; HUN-REN–UD Functional and Restoration Ecology Research Group, 1 Egyetem sqr., 4032 Debrecen, Hungary; Polish Academy of Sciences, Botanical Garden-Centre for Biological Diversity Conservation in Powsin, Warszawa, Poland

**Keywords:** Pannonian flora, persistence, plant functional groups, seed longevity, seed mass, seed morphology, seed shape, seed size, seed weight, soil seed bank

## Abstract

**Background and Aims:** Knowledge on seed persistence is vital from both theoretical and practical considerations but directly collecting seed persistence data for many species is rather unfeasible. Therefore, there is a need to identify traits that can predict seed persistence, but studies about the effects of seed size and shape on persistence yielded results varying across regions. We studied 392 species of the Pannonian flora (Central Europe) to asses (i) how seed mass and shape affect seed persistence, (ii) whether this effect is consistent across plant functional groups, and (iii) whether seed mass and shape are correlated in different functional groups?

**Methods:** We collected data on the seed mass and persistence of species and we performed measurements to calculate their Seed Shape Index, which quantifies deviation from sphericity. We analysed how seed mass and Seed Shape Index affect persistence in all herbaceous species and separately in four plant functional groups. We also tested whether and how seed mass and shape are related to each other in these groups.

**Key Results:** Across all species, both seed mass and Seed Shape Index negatively affected seed persistence. The same relationship was observed separately for perennials, short-lived species, and forbs, but only seed shape affected persistence in the graminoid group. Larger seeds also tended to be less spherical in graminoid species, in contrast to all studied species and to other functional groups, where we observed inverse or no relationship between seed mass and shape.

**Conclusions:** Consistent with many studies in other floras, both seed mass and shape negatively affected seed persistence in the Pannonian flora. However, only seed shape influenced persistence in graminoid species, suggesting that different factors may be at play in forbs and graminoids. Therefore, future studies of this relationship may need to treat and analyse graminoids separately.

## INTRODUCTION

Studying soil seed banks is vital for understanding the dynamics of plant populations and communities (Hopfensperger, 2007; Plue *et al*., 2017, 2020) and how different species deal with environmental heterogeneity and uncertainty (Long et al., 2015; Gioria *et al*., 2020). As soil seed banks disperse genetic diversity and mortality risks in time, they strongly promote the maintenance of plant populations (Gioria *et al*., 2020). Persistent soil seed bank formation decreases the risk related to reproductive failure during periods of adverse environmental conditions, thereby constituting a bet-hedging strategy (Venable and Brown, 1988). In plant communities exposed to environmental change, for example climate change or habitat isolation, persistent soil seed banks can decrease extinction risk and contribute to population persistence (Stöcklin and Fisher, 1999; Rees *et al*., 2002, Estrada *et al*., 2015). Soil seed banks also act as reserves of genetic variability (Levin, 1990; Aparicio *et al*., 2002). As persistent soil seed banks can contain seeds produced over multiple years and years with different environmental conditions benefit different genotypes of a species, the seed bank can provide a great diversity of seed genotypes adapted to varying environmental conditions (Cabin, 1996). Therefore, persistent soil seed banks play a vital role in the resilience of plant communities (Kiss *et al*., 2018) and the maintenance of biodiversity through space and time (Royo and Ristau, 2013). In spite of this, our knowledge on soil seed banks is still disproportionately limited compared to the aboveground vegetation.

For the formation of a persistent soil seed bank, seed persistence is a ubiquitous necessity. Seed persistence refers to the ability of seeds to remain viable for a long time (Fenner and Thompson, 2005), which allows them to wait in the soil seed bank until environmental conditions are right for their germination, thereby promoting survival under changing or unpredictable conditions (e.g., del Cacho and Lloret, 2012, Estrada *et al*., 2015). Seed persistence, which primarily varies on a continuous scale, is generally classified into discrete categories for simplicity and practical application. The most widespread seed bank classification system distinguishes three categories: (i) transient seeds (viable for <1 year), (ii) short-term persistent seeds (viable for 1–5 years), and (iii) long-term persistent seeds (viable for ≥5 years) (Thompson *et al*., 1997). However, the boundaries are not always clear, especially between the latter two categories (Thompson *et al*., 1993). Therefore, distinguishing between transient and persistent species – without differentiating between short-term and long-term persistence – is a reasonable approach for general discussions and broad analyses (see e.g., Bekker *et al*., 1998; Funes *et al*., 1999; Gioria *et al*., 2020). This distinction is especially important because it determines whether seeds of a species are able to accumulate over multiple seasons and therefore disperse in time (Baskin and Baskin, 2014).

The ability to distinguish species with transient versus persistent seeds is not only valuable for answering a wide range of fundamental questions in vegetation science and population biology (Saatkamp *et al*., 2009), but it is crucial for the success and feasibility of restoration projects as well (von Blanckenhagen and Poschlod, 2005; Török *et al*., 2018). Unfortunately, obtaining direct data on seed persistence is challenging and time-consuming (Cerabolini *et al*., 2003; Jaganathan *et al*., 2019). Soil seed bank analyses, whether using the seedling emergence or seed extraction method, require considerable effort. Moreover, the results are not necessarily conclusive, and the findings of different studies are frequently contradictory (Cerabolini *et al*., 2003). There are other methods to assess seed longevity more accurately, but they are either costly and challenging (such as carbon-dating viable seeds, e.g., Moriuchi *et al*., 2000), or take a particularly long time (such as burial experiments, e.g., Telewski and Zeevaart, 2002; Pakeman *et al*., 2012). Therefore, collecting direct seed persistence data for many species is rather unrealistic. Based on the above considerations, reliably predicting the ability of seeds to persist in the soil is the only realistic option for obtaining information on the transient versus persistent nature of a wide range of species. Since such information is highly needed from both theoretical and practical conservation perspectives (e.g., Saatkamp *et al*., 2009; Kalamees *et al*., 2012), investigating which attributes correlate with seed persistence and how these attributes can improve the reliability of predictions is vitally important.

Seed mass is the most frequently measured trait of seeds (Carta *et al*., 2024) and it is considered to have an exceptionally large functional importance, as it is connected to several processes and plant characteristics, such as dispersal distance, light detection, seedling establishment, seed predation, or the number of seeds produced (summarised for example by Moles, 2018 and Carta *et al*., 2024). Seed shape can also influence many processes such as soil penetration (Chambers *et al*., 1991), fire tolerance (Ruprecht *et al*., 2015), or the chance of surviving gut passage and therefore the potential for endozoochorous dispersal (van Leeuwen *et al*., 2023). However, it is more challenging to quantify and consequently studied less frequently (Dayrell *et al*., 2023; Carta *et al*., 2024).

Seed mass and shape have been repeatedly hypothesised to be related to seed persistence, with somewhat varying results. A connection between a persistent seed bank and small, spherical seeds has already been noted by Thompson in 1987. Subsequently, clear evidence for the correlation between the size, shape, and persistence of seeds has been demonstrated in the British flora by Thompson *et al*. (1993): all seeds were found to be persistent within a range defined by a maximum in seed mass and seed shape variance. Since then, this relationship has been studied in the flora of other regions as well. Several such studies have confirmed that persistent seeds tend to be smaller and more spherical than transient seeds in various floras, for example in Sweden (Bakker *et al*., 1996), Argentina (Funes *et al*., 1999), Iran (Thompson *et al*., 2001), Italy (Cerabolini *et al*., 2003), and China (Zhao *et al*., 2011). A recent study synthesising available data for 1,474 species worldwide also found that persistent seeds tend to be small and spherical (Wang *et al*., 2024). On the other hand, Peco *et al*. (2003), for example, found that although persistent seeds tend to be smaller, seed shape is not related to persistence in Mediterranean grasslands and scrublands in Spain. Conversely, McDonald *et al*. (1996) found that while persistent seeds did tend to be more spherical, seed weight was not related to persistence in a flood meadow in Great Britain. To complicate things even further, no relationship was detected between seed size, seed shape and persistence in some other regions, such as Australia (Leishman and Westoby, 1998) or South Africa (Holmes and Newton, 2004).

The relationship between seed traits and seed persistence appears to be highly context-dependent and varies across different floras worldwide. Both factors can be influenced by a range of environmental conditions (e.g., Harel *et al*., 2011; Abedi *et al*., 2014; Chen *et al*., 2021), while regional natural history and disturbance regimes further shape how variations in seed traits translate into differences in seed persistence (see e.g., Leishman and Westoby, 1998). Given these complexities, it is essential to assess the relationship between seed traits and seed persistence in various regions of the world. Expanding analyses to floras with varying climates and evolutionary histories will provide a more comprehensive understanding of the strength and generality of these relationships (Leishman and Westoby, 1998).

To our knowledge, how seed size and shape are related to seed persistence has never been studied in the Central European flora, presumably due to the scarcity of seed shape data for the species of Central Europe. To facilitate the analysis of the relationship between these factors, we set out to characterise the seed shape of a large number of plant species of the Pannonian Biogeographical Region, which is located in the eastern part of Central Europe and is bordered by the Carpathians, the Alps, and the Dinaric Mountains (EEA, 2016). We collected regional data on the seed mass and seed persistence of 392 species of the Pannonian flora and quantified the seed shape of all species. By analysing the compiled dataset, we aimed to answer the following questions: i) How do seed mass and seed shape influence seed persistence? ii) Is the effect of seed mass and shape on seed persistence consistent across plant functional groups? and iii) Are seed mass and shape related to each other and is this relationship consistent across plant functional groups?

## MATERIALS AND METHODS

### Data collection

To analyse the relationship of seed traits and seed persistence in the Pannonian flora, we collated a dataset fully based on regionally measured data, in order the avoid the potential confounding effects of distinct climates which can cause considerable intraspecific trait variability (Albert *et al*., 2010; Sonkoly and Török, 2024). The Pannonian Database of Plant Traits (PADAPT, Sonkoly *et al*., 2023) contains seed persistence data for more than 600 species based on regional soil seed bank studies. From these, we selected those species which are included in the seed collection of the Department of Ecology, University of Debrecen, and therefore available for seed shape measurements (424 species in total). In PADAPT, data on seed persistence is provided in the form of Seed Bank Persistence Index (SBPI) following the approach of the longevity index by Bekker *et al*. (1998). SBPI represents the proportion of data indicating the presence of a persistent seed bank for a species, ranging from zero to one. SBPI = 0 indicates that all available data suggest a transient seed bank for the species, while SBPI = 1 indicates that all available data suggest a persistent seed bank for the species. The thousand-seed mass (TSM) of the 424 species have already been measured on seeds stored in the aforementioned seed collection (see Török *et al*., 2013, 2016; Törő-Szijgyártó *et al*., 2023).

### Seed shape measurements

To quantify seed shape, we measured the width, length and thickness of the seeds of all 424 species. Twenty replicate measurements were performed for each species and then all width, length and thickness data were averaged between the 20 measurements. Thickness was measured using a HEDÜ 510-201 digital thickness gauge, with an accuracy of 0.02 mm. The length and width values of the same 20 seeds were obtained from photographs using WinImag 1.0 data acquisition system. In general, we aimed to measure diaspores in the form they are dispersed, meaning that the measured morphological unit was not necessarily a seed for all species, but here we refer to all of them as seeds for simplicity. Most grass and Asteraceae seeds were measured without appendage. *Rumex* seeds were also measured without appendage, as the presence of appendages makes accurate measurements difficult. The measured morphological unit for each species is given in Supplementary Table S1. The seed shape measurements were carried out on the same seed lots which were previously used for thousand-seed mass measurements (see Török *et al*., 2013, 2016; Törő-Szijgyártó *et al*., 2023), ensuring that the measured morphological units were the same for seed mass and shape measurements.

### Data analysis

To reduce the number of confounding factors in our analyses, we excluded two species groups from the analyses. We excluded aquatic plants because seed bank formation and seed persistence in aquatic habitats is presumably influenced differently by seed traits compared to terrestrial habitats. Following the approach of Powney *et al*. (2014), we categorised species with a soil moisture indicator value above 8 as aquatic (based on Borhidi, 1995) and excluded eight species from the analyses based on this criterium. Trees and shrubs were also excluded from the analyses as their seed persistence seems to be influenced by seed traits differently than that of herbaceous species (see Wang *et al*., 2024) and they were represented by too few species to be analysed separately. We categorised the species into life forms based on Sonkoly *et al*. (2023) and all phanaerophyte species including nanophanaerophytes (subshrubs), microphanerophytes (shrubs), and mega-mesophanerophytes (trees), altogether 24 species, were excluded from the analyses. After these exclusions, 392 species were included in the analyses (see Table S1).

As a measure of seed shape, we calculated the Seed Shape Index for each species. Seed Shape Index expresses how much the shape of a seed differs from being spherical, with a value of zero indicating a perfectly spherical seed. Increasing Seed Shape Index values indicate increasingly flattened and/or elongated seeds. For needle- or disc-shaped seeds, the maximum value is about 0.3 and varies very little between seeds of the same species. Following the calculations of Thompson *et al*. (1993), we calculated Seed Shape Index as the variance of seed length, width, and thickness. To prevent seed size from affecting the index, we first standardised the three dimensions by scaling them relative to seed length, which was set to 1.

To study the influence of TSM and Seed Shape Index on seed persistence, we used seed persistence as a binary dependent variable (transient vs. persistent). Following the approach of Gioria *et al*. (2020) and Wang *et al*. (2024) for example, Seed Bank Persistence Index (SBPI) values of zero were treated as having transient seeds, while species with an SBPI higher than zero were treated as having persistent seeds, because a SBPI higher than zero indicates that there was at least one study from Hungary finding a persistent seed bank for the species in question. We analysed the influence of TSM, Seed Shape Index, and their interaction on seed persistence as a binary variable (persistent vs. transient), using a logistic regression model. As a measure of effect size, we also calculated odds ratios. Because TSM and Seed Shape Index are on very different scales (TSM ranged from 0.004 g to 56.14 g, while Seed Shape Index ranged from 0.00013 to 0.29329), we used z-scoring prior to the analysis to standardise them so that both variables have a mean of zero and a standard deviation of one, which also makes the coefficients and odds ratios more comparable.

We also analysed the effect of TSM and Seed Shape Index on seed persistence separately for four plant functional groups: (i) forbs, (ii) graminoids, (iii) perennial species, and (iv) short-lived species. Life form was assigned to species according to Sonkoly *et al*. (2023) and we considered therophyte and hemitherophyte species to be short-lived (136 species). Species in other life form categories were considered perennial (256 species). Species in the Cyperaceae, Juncaceae and Poaceae families were considered graminoids (91 species), all other herbaceous species were considered to be forbs (301 species). We compared the TSM and the Seed Shape Index of persistent vs. transient species using Wilcoxon rank sum tests. We also tested the correlation between TSM and Seed Shape Index across all species and within the distinct functional groups, for which we used Spearman rank correlations because the data did not meet the assumptions of parametric tests. All analyses were carried out in an R environment (version 4.3.2, R Core Team, 2023). Nomenclature follows Euro+Med PlantBase (http://www.europlusmed.org).

## RESULTS

After the exclusion of aquatic and woody species, we performed the analyses with a dataset containing data on 392 species (see Table S1). In this dataset, TSM ranged from 0.004 g (*Gnaphalium uliginosum* and *Sagina procumbens*) to 56.14 g (*Iris pseudacorus*) while Seed Shape Index ranged from 0.00013 (*Vicia angustifolia*) to 0.29329 (*Stipa borysthenica*).

Across all 392 species, we found that both TSM and Seed Shape Index had a significant negative effect on seed persistence, with TSM having a stronger effect than Seed Shape Index (Table 1). The interaction term was also significant, indicating that at higher Seed Shape Index values the effect of TSM on persistence becomes less negative (see Fig. 1). The threshold TSM was found to be 0.054 g, meaning that all studied species with a thousand-seed mass lower than this have a persistent seed bank (Fig. 1). Wilcoxon rank sum tests showed that both the TSM (*p* < 0.00001, *W* = 8444) and the Seed Shape Index (*p* = 0.004, *W* = 11302) of transient species are significantly higher than those of persistent species, although the difference in Seed Shape Index is much less pronounced (Fig. 2).

**Table 1.** Results of the logistic regression model testing the effects of thousand-seed mass (TSM), Seed Shape Index, and their interaction on the seed bank persistence of all herbaceous species. TSM and Seed Shape Index values were scaled before the analysis (z-scoring). Significant differences are marked with italics.

|  | Coefficient<br>(Estimate) | Odds Ratio<br>(OR) | 95% Confidence<br>Interval (OR) | *p*-value |
| --- | --- | --- | --- | --- |
| TSM | -0.773 | 0.462 | 0.299–0.651 | *<0.001* |
| Seed Shape Index | -0.545 | 0.580 | 0.450–0.741 | *<0.0001* |
| TSM × Seed Shape Index | 0.434 | 1.544 | 1.136–2.142 | *0.007* |

**Figure 1.**
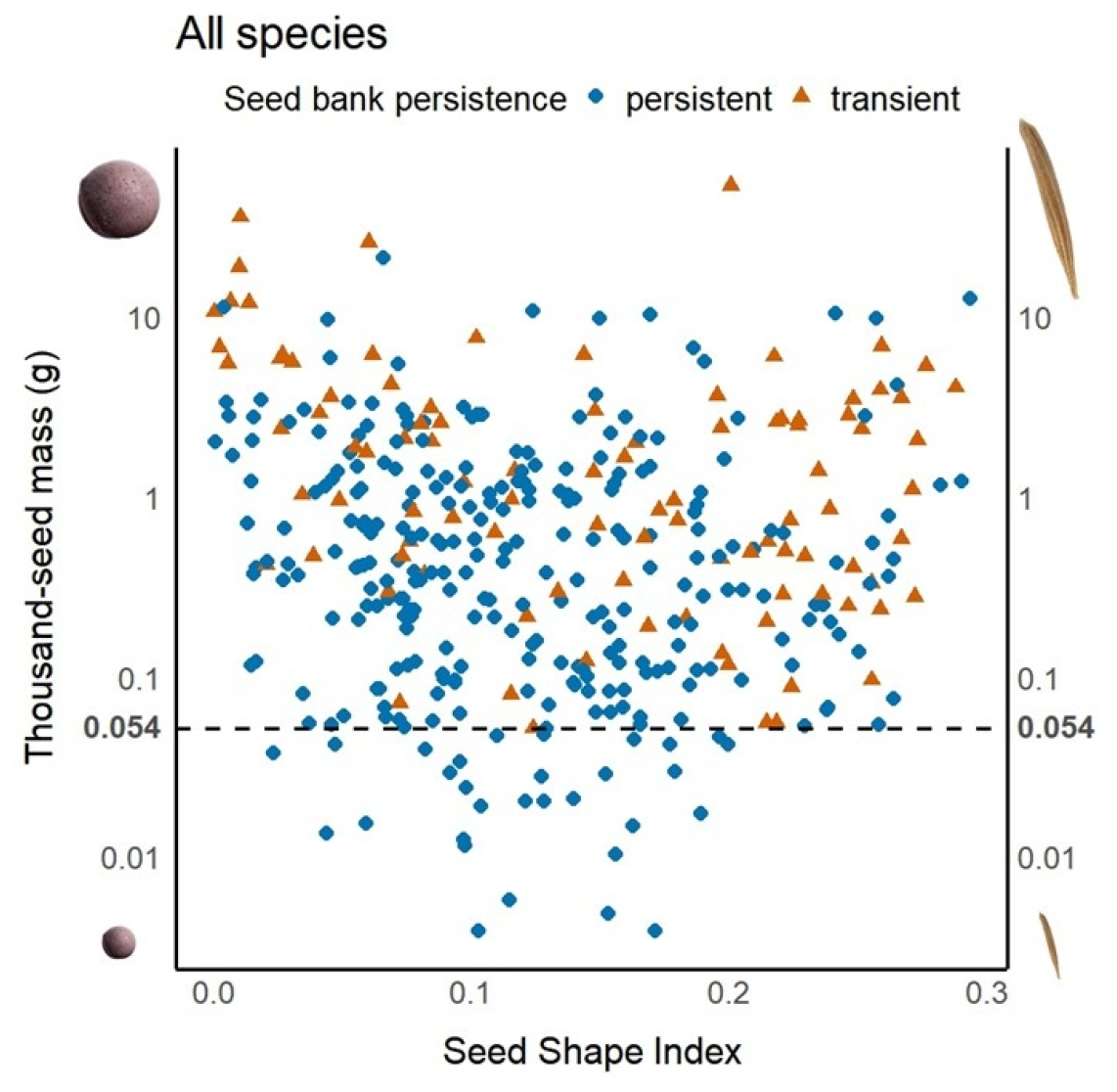
The relationship between seed mass, seed shape, and seed bank persistence in 392 herbaceous species of the Pannonian flora. The dashed line indicates the thousand-seed mass value (0.054 g) below which all studies species have persistent seed banks. Note that the y-axis is on a logarithmic scale.

**Figure 2.**
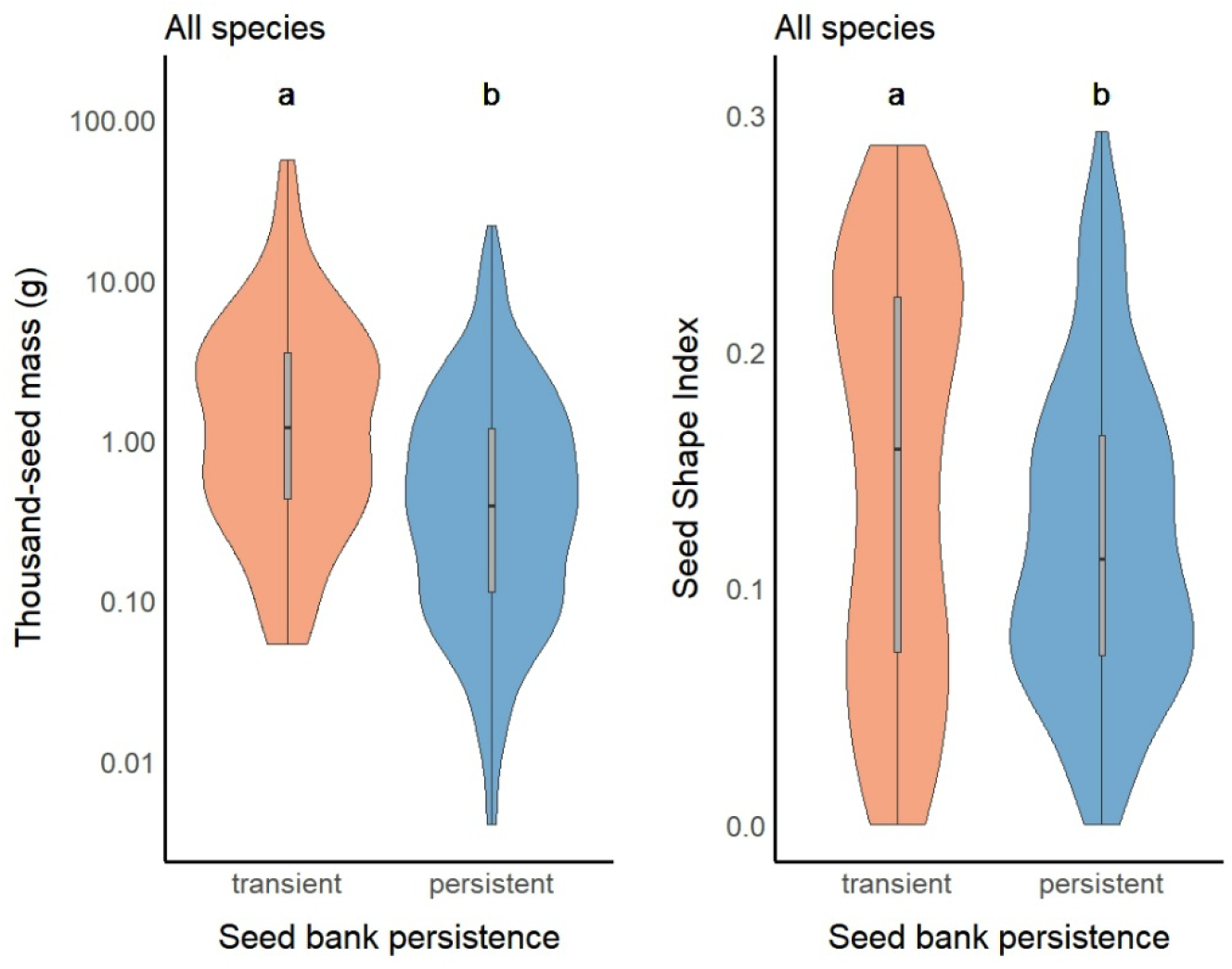
The thousand-seed mass (A) and Seed Shape Index (B) of species with transient vs. persistent seed banks. Different letters above the bars denote significant differences (Wilcoxon rank sum tests). Note that on figure A the y-axis is on a logarithmic scale.

To assess whether the effect of TSM and Seed Shape Index on seed persistence is consistent across functional groups, we studied the relationship separately in four functional groups. We found that in three of the four functional groups both TSM and Seed Shape Index significantly negatively affected seed persistence (Table 2, Fig. 3), which is consistent with the relationship across all species. On the other hand, only Seed Shape Index affected seed persistence in graminoid species, while TSM did not have a significant effect on it (Table 2). The interaction was significant in case of short-lived species (Table 2), indicating that at higher Seed Shape Index values the effect of TSM on persistence becomes less negative in short-lived species (see Fig. 3C), although the small number of species with transient seeds in the short-lived group might reduce the robustness of this analysis. Wilcoxon rank sum tests showed that the TSM of transient species was significantly higher than that of persistent species in all functional groups, but a significant difference in Seed Shape Index was only detected in graminoid species (for the detailed results see Supplementary Table S2 and Fig. S1).

**Table 2.** Results of logistic regression models testing the effects of thousand-seed mass (TSM), Seed Shape Index, and their interaction on the seed bank persistence of species in different functional groups. TSM and Seed Shape Index values were scaled before the analysis (z-scoring). Significant differences are marked with italics.

|  | **Coefficient**<br>**(Estimate)** | **Odds Ratio**<br>**(OR)** | **95% Confidence**<br>**Interval (OR)** | ***p*-value** |
| --- | --- | --- | --- | --- |
| **Forb species (n=301)** |  |  |  |  |
| TSM | -0.893 | 0.410 | 0.226–0.613 | *<0.001* |
| Seed Shape Index | -0.359 | 0.699 | 0.526–0.925 | *0.012* |
| TSM × Seed Shape Index | 0.393 | 1.481 | 0.910–2.164 | 0.063 |
| **Graminoid species (n=91)** |  |  |  |  |
| TSM | -0.423 | 0.656 | 0.295–1.501 | 0.296 |
| Seed Shape Index | -0.960 | 0.383 | 0.202–0.665 | *0.001* |
| TSM × Seed Shape Index | 0.471 | 1.601 | 0.838–3.217 | 0.157 |
| **Perennial species (n=256)** |  |  |  |  |
| TSM | -0.931 | 0.394 | 0.192–0.680 | *0.006* |
| Seed Shape Index | -0.394 | 0.674 | 0.506–0.892 | *0.006* |
| TSM × Seed Shape Index | 0.408 | 1.504 | 0.927–2.516 | 0.098 |
| **Short-lived species (n=136)** |  |  |  |  |
| TSM | -0.842 | 0.431 | 0.235–0.749 | *0.004* |
| Seed Shape Index | -0.889 | 0.411 | 0.213–0.746 | *0.005* |
| TSM × Seed Shape Index | 0.460 | 1.585 | 1.053–2.524 | *0.034* |

**Figure 3.**
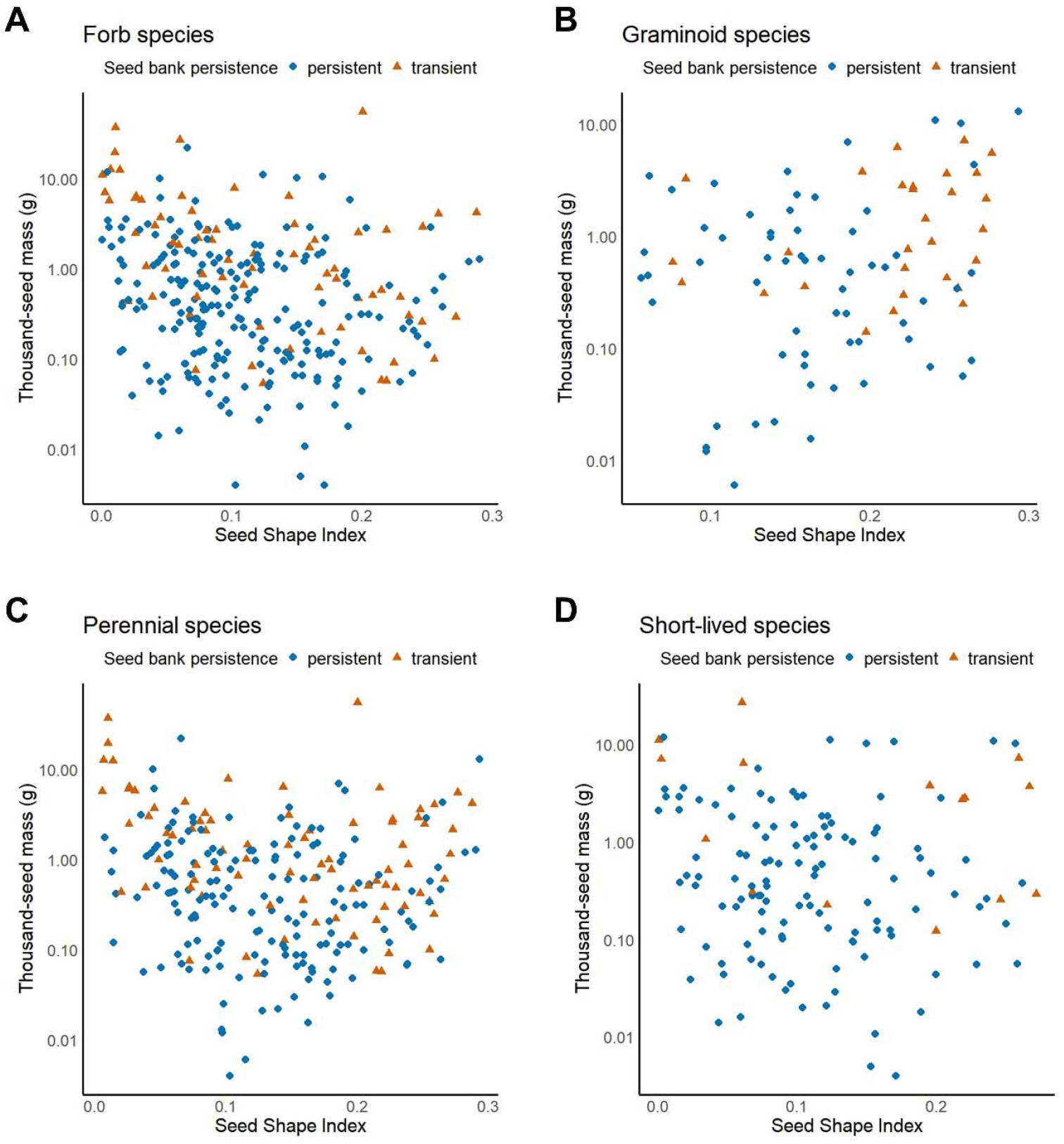
The relationship between seed mass, seed shape, and seed bank persistence in different functional groups: A – forb species, B – graminoid species, C – perennial species, D – short-lived species.

Across all species, we found a weak negative correlation between TSM and Seed Shape Index (Fig. 4A). Similar negative correlations were found between these two variables in the forb and perennial functional groups (Fig. 4B and 4D), while there was no significant correlation in the case of short-lived species (Fig. 4E). The relationship was positive in the case of graminoid species (Fig. 4C).

**Figure 4.**
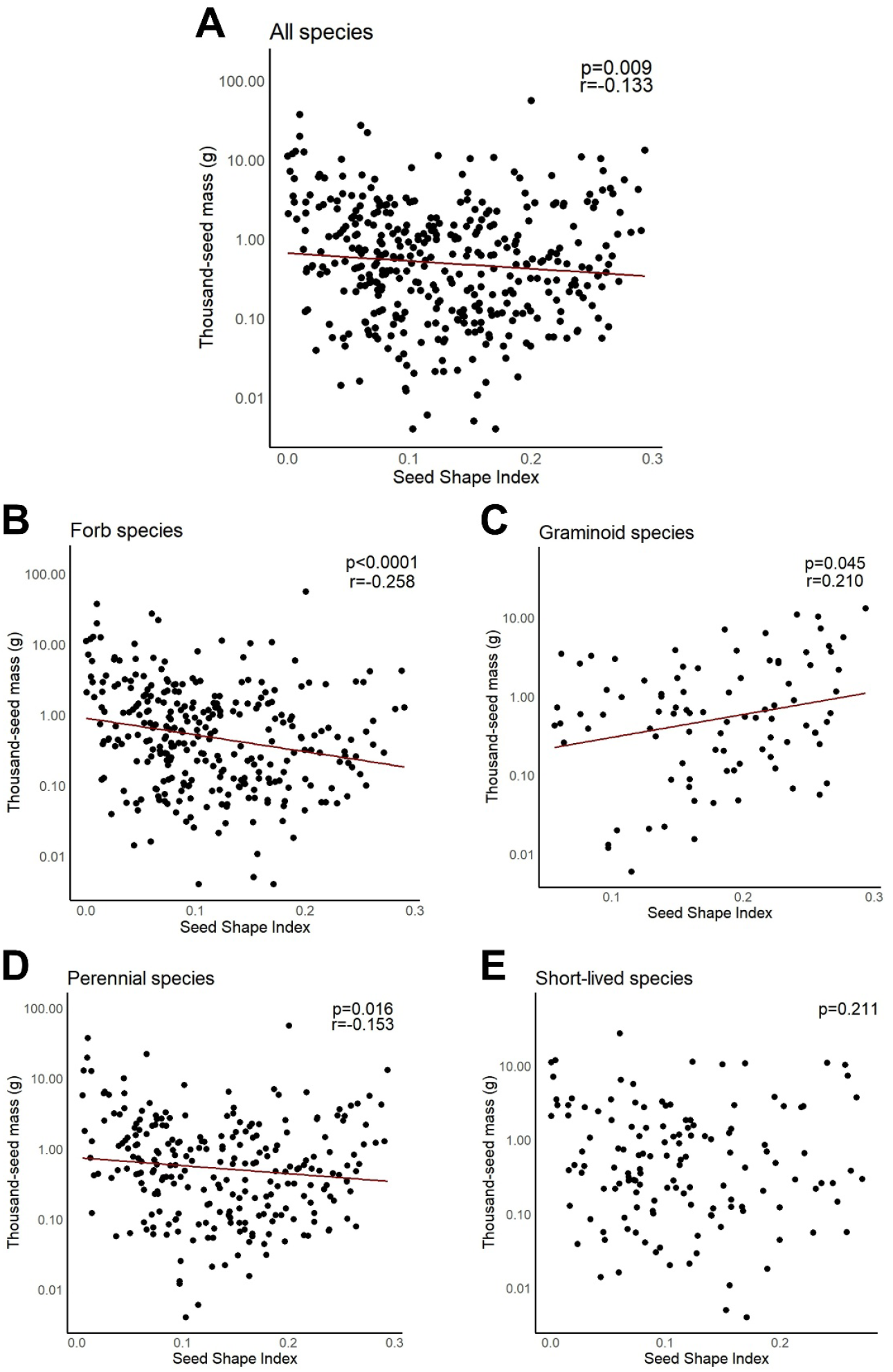
Correlation between seed mass and seed shape in all the 392 herbaceous species (A) and in different functional groups: A – forb species, B – graminoid species, C – perennial species, D – short-lived species. Note that the y-axes are on a logarithmic scale.

## DISCUSSION

In line with the findings of several previous studies (e.g., Thompson *et al*., 1993; Funes *et al*., 1999; Zhao *et al*., 2011), we found that both seed mass and seed shape are significantly negatively related to seed persistence in 392 herbaceous species of the Pannonian flora. However, seed shape appears to be a less important driver of seed persistence in this region. Several previous studies have found that only seed size is significantly related to seed persistence in various regions of the world (e.g., Bekker *et al*., 1998; Peco *et al*., 2003; Yu *et al*., 2007; Wang *et al*., 2011). Therefore, it seems to be a rather common trend that seed shape is less important than seed size in determining seed persistence. However, there are also results implying that only seed shape significantly affects seed persistence (McDonald *et al*., 1996).

Seeds with a thousand-seed mass below 0.054 g were all persistent, implying that in the Pannonian flora all seeds with a mass below this threshold value may be considered persistent with a reasonable certainty. However, many persistent seeds were relatively large, therefore, an upper threshold, above which all seeds could be considered transient, cannot be identified. Similarly, although spherical seeds were found to be more likely to be persistent, many persistent seeds were markedly non-spherical, in line with the findings of Moles *et al*. (2000).

Our results agree with the notion that seed persistence cannot be reliably predicted based only on seed mass (Gioria *et al*., 2020), seed shape or perhaps other seed characteristics such as seed coat thickness, dormancy mechanisms or nutrient reserves may also need to be considered (e.g., Davis *et al*., 2016; Zalamea *et al*., 2018). Using seed bank persistence data provided by Thompson *et al*. (1997) and seed mass data measured in the Pannonian region, Csontos and Tamás (2003) demonstrated that the proportion of transient species is increasing with increasing seed mass in the Pannonian flora. This indicates that the relationship between seed size and persistence is the same in the Pannonian flora as in most other regions where it was studied, but seed shape has not been considered in this analysis. Moreover, the analysed seed bank persistence data originated from different regions, which may have a confounding effect.

Although Wang *et al*. (2024) found no interaction between seed mass and seed shape, the interaction between them was significant in our analysis encompassing all species. However, in contrast to the analysis of Wang *et al*. (2024), our analysis only included herbaceous non-aquatic species. The negative interaction term indicated that at higher Seed Shape Index values the effect of TSM on persistence is less negative, which may be the result of the fact that above a Seed Shape Index of approximately 0.2, there were no species with very small seeds in the dataset. This negative interaction may not exist in floras containing several species with very small, non-spherical seeds.

Seed mass and shape may be related to seed persistence due to a number of reasons. The size and shape of seeds presumably affect their ability to move towards deeper soil layers (e.g., Bekker *et al*., 1998; Schmiede *et al*., 2009). For example, there are studies suggesting that large and elongated seeds are less likely to be buried by the activity of soil biota (Thompson *et al*., 1994; Bernhardt, 1995). One theory is that the correlation may be due to this tendency of small and isodiametric seeds to quickly become buried in the soil, because being able to persist until a disturbance brings them to the soil surface again may commonly be necessary for these seeds (Moles *et al*., 2000). As seeds experience higher rates of predation on the soil surface compared to when they are buried (Hulme, 1998; Jacob *et al*., 2006), it can also be hypothesised that large and elongated or flattened seeds with slow soil penetration cannot escape predation by quickly being buried in the soil (Hulme and Borelli, 1999). Therefore, for these species it may be less advantageous to build a persistent seed bank. Buried seeds are also less exposed to germination-stimulating temperature fluctuations and light (Fenner and Thompson, 2005). Larger seeds are generally able to germinate from deeper soil layers (Grundy *et al*., 2003; Sonkoly *et al*., 2020), while soil burial typically hinders the germination of small seeds, because they are more likely to have a light requirement for germination (Milberg *et al*., 2000). This means that they are more likely to remain ungerminated once they are buried, providing them an opportunity to persist in the soil.

Whether the relationship between seed persistence and the size and shape of seeds varies between different plant functional groups has also not been studied yet. For example, it is known that to ensure survival during periods of unfavourable environmental conditions, short-lived species more strongly depend on persistent seeds than perennial species (Meyer *et al*., 2006; Scott *et al*., 2010). Accordingly, short-lived plant species tend to have more persistent seeds than perennials (Gioria *et al*., 2020), which are typically also smaller-sized than the seeds of perennial species (Thompson *et al*., 1998; Wang *et al*., 2011). A short life-span can also be associated with persistent seeds through the disturbance regime of the habitat. The proportion of short-lived species is higher in disturbed habitats than in relatively undisturbed ones, as disturbance can lead to changes in plant community composition in favour of species with rapid growth and with a resource-acquisitive strategy (Smith *et al*., 2022). As ensuring the survival of the population in disturbed habitats requires the formation of a persistent seed bank (Fenner and Thompson, 2005), the relationship between seed traits and seed persistence may not be the same in short-lived and perennial species. In this context, it has already been demonstrated that different plant functional groups such as annuals and perennials can have contrasting relationships between seed size and several other factors like competitive ability and seed production (Coomes and Grubb, 2003). Seed bank types can also be contrasting in forb and graminoid species even within the same habitat type (Bertiller and Aiola, 1997), and the relationship between habitat characteristics and the proportion of species with persistent seeds can differ significantly in forb and graminoid species (Zeiter *et al*., 2013). Moreover, the ability of a species’ seeds to persist is also related to phylogeny (Gioria *et al*., 2020).

Based on the above considerations, the influence of seed traits on seed persistence may vary considerably across different plant functional groups, and our findings confirm this assumption. We found that both seed mass and seed shape affect seed persistence analysed across all species and in the forb, perennial, and short-lived groups, with seed mass having a stronger effect in all the above groups except for the short-lived one. In contrast to this, only seed shape significantly influenced seed persistence in the case of graminoid species. These results suggest that there are different effects at play in forb and in graminoid species. Wang *et al*. (2024) found that the relationship of seed mass and shape with seed persistence is not consistent across phylogenetic clades. According to their findings, both seed mass and shape affect persistence in Poales, but only seed mass affects persistence in Asterales and Lamiales, while no significant effect was detected in Fabales and Caryophyllales. Although the nature of the relationship they found in Poales is not the same as what we found in the graminoid group (which consisted mainly of Poales species), their findings are consistent with our results in the sense that the relationship between seed traits and seed persistence in graminoids is different from the relationship seen in other species. Moreover, the relationship of these variables may also differ between short-lived and perennial graminoid species. Taken together, these findings suggest that graminoids exhibit distinct relationships and might have to be treated and analysed separately from other species to disentangle the complex relationships between seed traits and seed bank persistence.

If the effect of seed size and shape on seed persistence is not consistent across different plant functional groups, it may be because they are differently related to each other in different functional groups, which could at least partially explain inconsistencies. However, most previous studies about the influence of seed size and shape on seed persistence have not assessed whether and how seed size and shape themselves are correlated (e.g., Moles *et al*., 2000; Peco *et al*., 2003; Wang *et al*., 2024), leaving this question unresolved. To our knowledge, the study of Zhao *et al*. (2011) is the only exception. They studied 141 species of sand grasslands in Northern China and found a slight tendency of bigger seeds to be more spherical, but the relationship was not significant.

In this study, there was a weak negative correlation between seed mass and Seed Shape Index in all species and in the functional groups of perennials and forbs, while there was no significant correlation in short-lived species. On the other hand, there was a weak but significant positive correlation between the two variables in graminoid species. Therefore, in graminoids, seed shape may counteract or modify the commonly observed effect of seed mass on persistence, making seed mass a non-significant predictor of persistence in this group. In the studied 91 graminoid species of the Pannonian flora, small-seeded species were generally found to have more spherical seeds (e.g., *Juncus* or *Agrostis* species), while larger graminoid seeds are typically more elongated (e.g., *Stipa* or *Bromus* species). Csontos and Kalapos (2013) also found that larger seeds tend to be less isodiametric in 137 grass species of the Pannonian flora with C3 photosynthesis. This trend therefore seems to be quite obvious in the Pannonian flora, but this might not be the case in other regions of the world, which could also cause a different relationship between seed size and shape and seed persistence in the graminoids of other floras. A possible future direction would therefore be to test whether the seed persistence of graminoid species is also influenced solely by seed shape in other floras and whether the negative association between seed size and seed sphericity observed in the graminoids of the Pannonian flora exists in other floras as well.

## CONCLUSIONS

Persistent seed banks have a key role in community resilience and in the maintenance of biodiversity through space and time; therefore, enhancing our knowledge of the formation of persistent seed bank is vitally important. Our analysis represents an advance in our ability to successfully predict seed persistence in the soil. We found that similarly to the floras of several other regions, both seed size and seed shape are significantly related to seed persistence in the Pannonian flora, with seed shape having a less strong influence. By analysing the relationship between these factors in different plant functional groups separately, we also revealed that graminoid species show distinct relationships. Therefore, although the general trend found in our study is consistent with most previous analyses of seed size, seed shape, and persistence, our more detailed results regarding different plant functional groups suggest that detailed analyses are necessary in the floras of other regions as well and future studies of this relationship may need to treat and analyse graminoid species separately.

## SUPPLEMENTARY INFORMATION

**Table S1:** The generated dataset containing the Seed Bank Persistence Index, thousand-seed mass, seed length, seed width, seed thickness, Seed Shape Index, and further information about the studied 392 species of the Pannonian flora.

**Table S2:** Results of the Wilcoxon rank sum tests comparing transient and persistent species in terms of thousand-seed mass and Seed Shape Index separately in different plant functional groups.

**Figure S1.** The difference between transient and persistent species in terms of thousand-seed mass and Seed Shape Index in different plant functional groups.

## AUTHOR CONTRIBUTIONS

V.T-S: investigation, methodology, data curation, writing–original draft. P.T: conceptualization, methodology, funding acquisition, writing–review & editing. K.T: investigation. H.M-R: investigation. L.R.G.S: investigation. S.M: investigation. G.K-V: investigation, writing–review & editing. A.M-B: investigation. P.D.C: investigation. J.S: conceptualization, data curation, formal analysis, funding acquisition, writing–original draft.

## ACKNOWLEDGEMENTS

We are grateful to the many colleagues who provided seeds for the seed collection used for measurements.

## DATA AVAILABILITY STATEMENT

All the data generated for and used in this study is available in Supplementary Table S1.

