## Supplementary Information (Figure S1 and Table S2) for "Inconsistent relationships detected between seed size, shape, and persistence for different plant functional groups in the Pannonian flora"

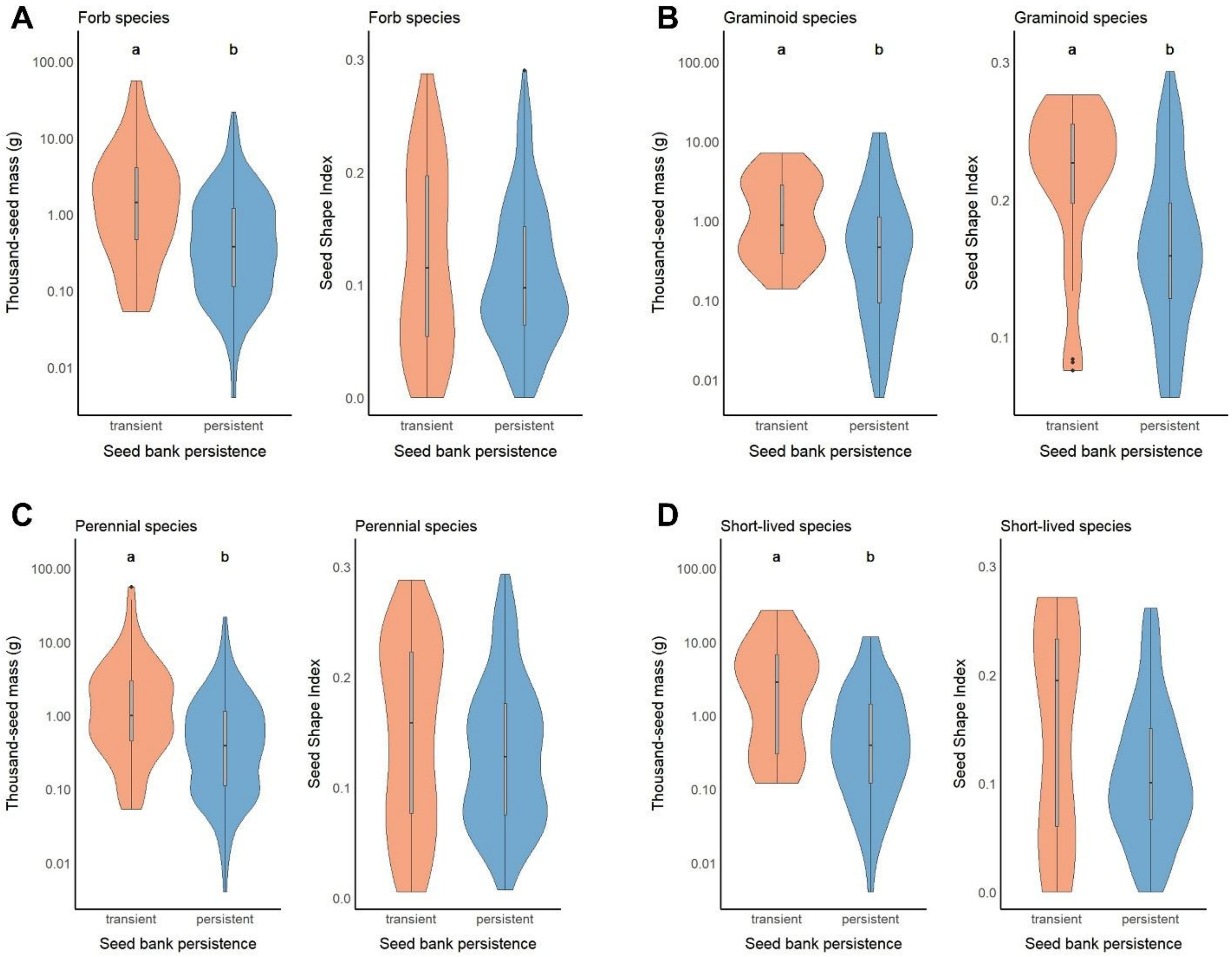


**Figure S1.** The thousand-seed mass and Seed Shape Index of transient and persistent species in different plant functional groups of 392 herbaceous species of the Pannonian flora. Significant differences based on Wilcoxon rank sum tests are indicated by different letters above the bars. A – forb species, B – graminoid species, C – perennial species, D – short-lived species.

**Table S2.** Results of the Wilcoxon rank sum tests comparing transient and persistent species in terms of thousand-seed mass (TSM) and Seed Shape Index separately in different plant functional groups. Significant differences are marked with italics.

| | ***W*** | ***p*** |
|---|---|---|
| **Forb species (n=301)** | | |
| TSM | 4632 | <*0.001* |
| Seed Shape Index | 8317 | 0.402 |
| **Graminoid species (n=91)** | | |
| TSM | 586.5 | *0.008* |
| Seed Shape Index | 1335 | <*0.001* |
| **Perennial species (n=256)** | | |
| TSM | 4109 | <*0.001* |
| Seed Shape Index | 7810 | 0.071 |
| **Short-lived species (n=136)** | | |
| TSM | 476 | *0.003* |
| Seed Shape Index | 1101 | 0.180 |
